# Consolidated bioprocessing of corn cob-derived hemicellulose: engineered industrial *Saccharomyces cerevisiae* as efficient whole cell biocatalysts

**DOI:** 10.1101/2020.07.01.182345

**Authors:** Joana T. Cunha, Aloia Romaní, Kentaro Inokuma, Björn Johansson, Tomohisa Hasunuma, Akihiko Kondo, Lucília Domingues

## Abstract

Consolidated bioprocessing, which combines saccharolytic and fermentative abilities in a single microorganism, is receiving increased attention to decrease environmental and economic costs in lignocellulosic biorefineries. Nevertheless, the economic viability of lignocellulosic ethanol is also dependent of an efficient utilization of the hemicellulosic fraction, which is mainly composed of xylose and may comprise up to 40 % of the total biomass. This major bottleneck is mainly due to the necessity of chemical/enzymatic treatments to hydrolyze hemicellulose into fermentable sugars and to the fact that xylose is not readily consumed by *Saccharomyces cerevisiae* – the most used organism for large-scale ethanol production. In this work, industrial *S. cerevisiae* strains, presenting robust traits such as thermotolerance and improved resistance to inhibitors, were evaluated as hosts for the cell-surface display of hemicellulolytic enzymes and optimized xylose assimilation, aiming at the development of whole-cell biocatalysts for consolidated bioprocessing of corn cob-derived hemicellulose. These modifications allowed the direct production of ethanol from non-detoxified hemicellulosic liquor obtained by hydrothermal pretreatment of corn cob, reaching an ethanol titer of 11.1 g/L corresponding to a yield of 0.328 gram per gram of potential xylose and glucose, without the need for external hydrolytic catalysts. Also, consolidated bioprocessing of pretreated corn cob was found to be more efficient for hemicellulosic ethanol production than simultaneous saccharification and fermentation with addition of commercial hemicellulases. These results show the potential of industrial *S. cerevisiae* strains for the design of whole-cell biocatalysts and paves the way for the development of more efficient consolidated bioprocesses for lignocellulosic biomass valorization, further decreasing environmental and economic costs.

## Introduction

Lignocellulosic biomass is a sustainable resource for the production of biofuels, and its utilization is a possible solution to alleviate the current world dependence on fossil fuels. In this context, consolidated bioprocessing (CBP), in which the same microorganism is able to produce hydrolytic enzymes and ferment sugars into ethanol [1], emerges as a promising alternative configuration to simultaneous saccharification and fermentation (SSF) process [2] and holds great promise for the efficient conversion and valorization of lignocellulosic biomass. Enzymes can be either secreted or displayed on the cell surface. The cell-surface display strategy exhibits several advantages, such as: i) high localized enzyme activity, ii) the monosaccharide release occurs close to the cell surface being instantaneously consumed by the cell, which reduces the risk of contamination or product inhibition, iii) the immobilization of enzymes on cell surface allows their re-utilization in successive cultures, which lowers the overall process cost [3]. Moreover, this strategy enables the use of recombinant microorganisms as whole-cell biocatalysts. The whole-cell biocatalysis provides a sustainable alternative to traditional chemical catalysis, since biocatalysis presents higher selectivity and catalytic efficiency, can be carried out at milder operation conditions and multi-step reactions can be performed in a single strain allowing cofactor regeneration [4]. Moreover, the CBP strategy could contribute to a lower bioethanol production cost from lignocellulosic biomass, since CBP combines three process stages in a single recombinant microorganism, namely: the enzyme production, enzymatic saccharification and sugar fermentation. The effective use of hemicellulose is fundamental for bioethanol production as xylan may be a major constituent of some lignocellulosic biomasses, e.g. corn cob can achieve up to 31 g of xylan per 100 g of raw material [5]. Xylan from the hemicellulosic fraction of agro-industrial residues is an amorphous heteropolymer that comprises a backbone of β-1,4-linked xylose partially substituted with acetyl groups, uronic acids and arabinose [6]. The main chain of xylan can be hydrolysed into xylooligosaccharides (XOS) by endoxylanase, which can be cleaved into xylose by xylosidase [7]. Hydrothermal treatment of lignocellulosic biomass is a recognized environmental friendly process that improves the enzymatic saccharification of cellulose promoting the solubilization of hemicellulose as hemicellulosic-derived compounds (mainly composed by XOS). Still, these XOS are not metabolized by the yeast *Saccharomyces cerevisiae*, main microbial ethanol producer. Thus, the direct conversion of hemicellulose into ethanol requires the simultaneous expression of xylan-degrading enzymes and xylose-assimilating enzymes in the recombinant S. *cerevisiae* strain. Despite promising advantages of CBP process over other alternatives, few attempts have been reported for the direct conversion of hemicellulose into ethanol, achieving ethanol concentrations in the range from 0.32 to 8.2 g/L [6, 8-12]. The most remarkable results with cell-surface display in a hemicellulosic CBP were obtained with liquors derived from hydrothermal treatment of rice straw, in which 4.04 and 8.2 g/L of ethanol were produced by engineered industrial *S. cerevisiae* Sun049 strain and engineered laboratorial *S. cerevisiae* NBRC1440/X strain, respectively [6, 8].

One of the most important limitation of CBP process is the selection of operating temperature, since ethanologenic yeast (such as *Saccharomyces cerevisae*) and xylanases/cellulases work at different range of temperatures. In addition, the engineered yeast has to function under adverse conditions, such as the presence of inhibitors derived from pretreatment (including acetic acid, furfural and 5-hydroxymethylfurfural (HMF)). In this sense, the selection of host microorganism with desired features such as thermotolerance and resistance to inhibitors is crucial to develop this sustainable bioprocess. Previous works have demonstrated the specific robustness [13, 14] and superior capacity of yeast strains isolated from industrial environments for lignocellulosic fermentation in these demanding conditions [15]. In addition, hemicellulosic CBP requires a highly engineered yeast, as xylose consumption pathways must be expressed together with xylan degrading enzymes. In this context, recent works have indicated that industrial isolates can present different intrinsic abilities to cope with genetic engineering strategies for xylose consumption [16] and expression of hydrolases [17, 18]. Therefore, there is a need for a tailor-made development of hemicellulosic ethanol-producing yeast where intrinsic capabilities of host microorganism (suitable for the process) are previously selected.

Taking into account the interest of the development of CBP as a green alternative to chemical catalysis in a lignocellulose biorefinery context, the aim of this study was to explore the robustness features of industrial *S. cerevisiae* isolates for cell-surface display of xylan-degrading enzymes together with the expression of xylose consumption pathways, as well as, to evaluate their performance in non-detoxified corn cob liquor as whole-cell biocatalysts for hemicellulose saccharification and direct conversion into ethanol.

## Results and Discussion

### Corn cob processing: hydrothermal treatment for hemicellulosic liquors

In order to evaluate the capacity of the constructed strains for the enzymatic saccharification and fermentation of hemicellulose, corn cob was selected as a representative renewable resource due to its high xylan content. The chemical composition of corn cob (expressed in g/100 g of raw material in oven-dry ± standard deviation based on three replicate determinations) was as follows: 28.79 ± 1.45 of glucan, 29.63 ± 0.45 of xylan, 3.62 ± 0.13 of arabinan, 2.58 ± 0.05 of acetyl groups and 18.58 ± 0.87 of Klason lignin. Corn cob was submitted to hydrothermal treatment under conditions selected based on previous works [5, 19]. Table 1 shows the chemical composition of solid and liquid phases after pretreatment at different severities (S_0_, 3.67, 3.99 and 3.78, corresponding to Tmax of 205, 211 and 207 °C in non-isothermal regime) and liquid solid ratios (LSR, 4 g/g and 8 g/g). The resulting hemicellulosic liquors (liquid phase) were denominated according to their potential xylose (g/L), i.e. Liquor 29X_Pot_, 32X_Pot_ and 54X_Pot_, accordingly. Solid phase was composed mainly by cellulose (measured as glucan) and lignin. As seen in Table 1, 81-89 % of xylan was solubilized during the pretreatment. XOS were the major component in the liquid phase, corresponding to 65-67 % of total identified compounds. In the treatments at LSR of 8 g/g, similar concentration of XOS were obtained at the two severities evaluated (3.67 and 3.99). Nevertheless, maximal recovery of xylan as sum of XOS and xylose (32 g/L) was obtained at severity of 3.99 (Liquor 32X_Pot_). Moreover, glucooligosaccharides were also quantified achieving a concentration in the range of 1.86-3.36 g/L (Table 1). Regarding degradation compounds (such as furfural and hydroxymethylfurfural-HMF), their concentration was higher at severity of 3.99 than at S_0_=3.67. In order to evaluate the capacity of strains at different substrate concentrations, liquid to solid ratio in the hydrothermal pretreatment was reduced to 4 g/g to obtain a liquor with higher XOS and xylose concentration, achieving a xylose potential (measured as sum of XOS and xylose) of 54 g/L (Liquor 54X_Pot_). Nevertheless, undesired compounds, including acetic acid, furfural and HMF, were also increased in this hemicellulosic liquor, even with the decrease of the treatment severity to 3.79 (Table 1), which may have an inhibitory effect on yeast growth [20].

**Table 1.** Composition of the pretreated corn cob and hemicellulosic liquors resultant of different hydrothermal treatments.

| **Pretreatment conditions** | **S<sub>0</sub>=3.67 and LSR=8 g/g** | **S<sub>0</sub>=3.99 and LSR=8 g/g** | **S<sub>0</sub>=3.78 and LSR=4 g/g** |
| --- | --- | --- | --- |
| Solid yield (g of pretreated corn cob/100 g of corn cob) | 57 | 42 | 57 |
| *Pretreated corn cob composition (g of component/100 g of pretreated corn cob)* |  |  |  |
| Glucan | 49.6 ± 0.60 | 63.32 ± 2.20 | 54.46 ± 1.59 |
| Xylan | 10.05 ± 0.10 | 5.74 ± 0.12 | 10.80 ± 0.30 |
| Arabinan | 1.21 ± 0.10 | 0.39 ± 0.02 | 0.66 ± 0.02 |
| Acetyl groups | 0.53 ± 0.01 | 0.52 ± 0.06 | 0.87 ± 0.07 |
| Klason Lignin | 19.4 ± 0.60 | 18.53 ± 0.56 | 21.58 ± 0.88 |
| *Liquid phase composition (g/L)* |  |  |  |
| **Liquors** | **29X<sub>Pot</sub>** | **32X<sub>Pot</sub>** | **54X<sub>Pot</sub>** |
| Glucose | 0.435 ± 0.012 | 0.353 ± 0.055 | 0.547 ± 0.023 |
| Xylose | 2.00 ± 0.07 | 5.01 ± 0.02 | 5.49 ± 0.22 |
| Arabinose | 1.46 ± 0.04 | 1.33 ± 0.02 | 2.37 ± 0.10 |
| Acetic acid | 1.06 ± 0.04 | 1.59 ± 0.01 | 2.42 ± 0.08 |
| Hydroxymethylfurfural (HMF) | 0.0734 ± 0.0024 | 0.123 ± 0.000 | 0.166 ± 0.007 |
| Furfural | 0.408 ± 0.011 | 0.649 ± 0.000 | 1.40 ± 0.04 |
| Glucoligosaccharides | 2.04 ± 0.06 | 1.86 ± 0.06 | 3.36 ± 0.04 |
| Xylooligosaccharides | 26.5 ± 0.9 | 26.7 ± 0.9 | 48.8 ± 1.4 |
| Arabinooligosaccharides | 1.63 ± 0.19 | 1.25 ± 0.06 | 2.43 ± 0.23 |
| Acetyl groups | 2.29 ± 0.11 | 2.25 ± 0.14 | 4.6 ± 0.34 |

Treatment were performed under non-isothermal conditions. S<sub>0</sub>: severity. LSR: liquid to solid ratio.

### Evaluation of engineered yeast strains’ hydrolytic capacity on hemicellulose derived compounds

*Aspergillus aculeatus* β-glucosidase 1 (BGL1), *Aspergillus oryzae* β-xylosidase A (XYLA) and *Trichoderma reesei* endoxylanase II (XYN) were displayed in the cell surface of the commercial bioethanol strain Ethanol Red, of the isolates from first generation bioethanol plants in Brazil, PE-2 and CAT-1 strains, and of an isolate from “cachaça” fermentation from Brazil, CA11. The BGL1 enzyme was additionally expressed to saccharify the glucooligosaccharides (GOS) also present in the hemicellulosic liquors (Table 1). Then, the capacity of the different *S. cerevisiae* industrial strains as whole cell biocatalysts for the saccharification of two hemicellulose derived compounds with different polymerization degrees (commercial beechwood xylan and hemicellulosic liquor obtained from hydrothermal treatment of corncob as described above) was evaluated. The modified strains, ER-X, PE-2-X, CAT-1-X and CA11-X, were characterized in terms of xylanase activity at 30 and 40 °C (Table S1 [see Additional file 1]) and capacity of saccharification of xylan from beechwood and corn cob hemicellulosic liquor 29X_Pot_ (Fig. 1, Table 2).

**Table 2.**
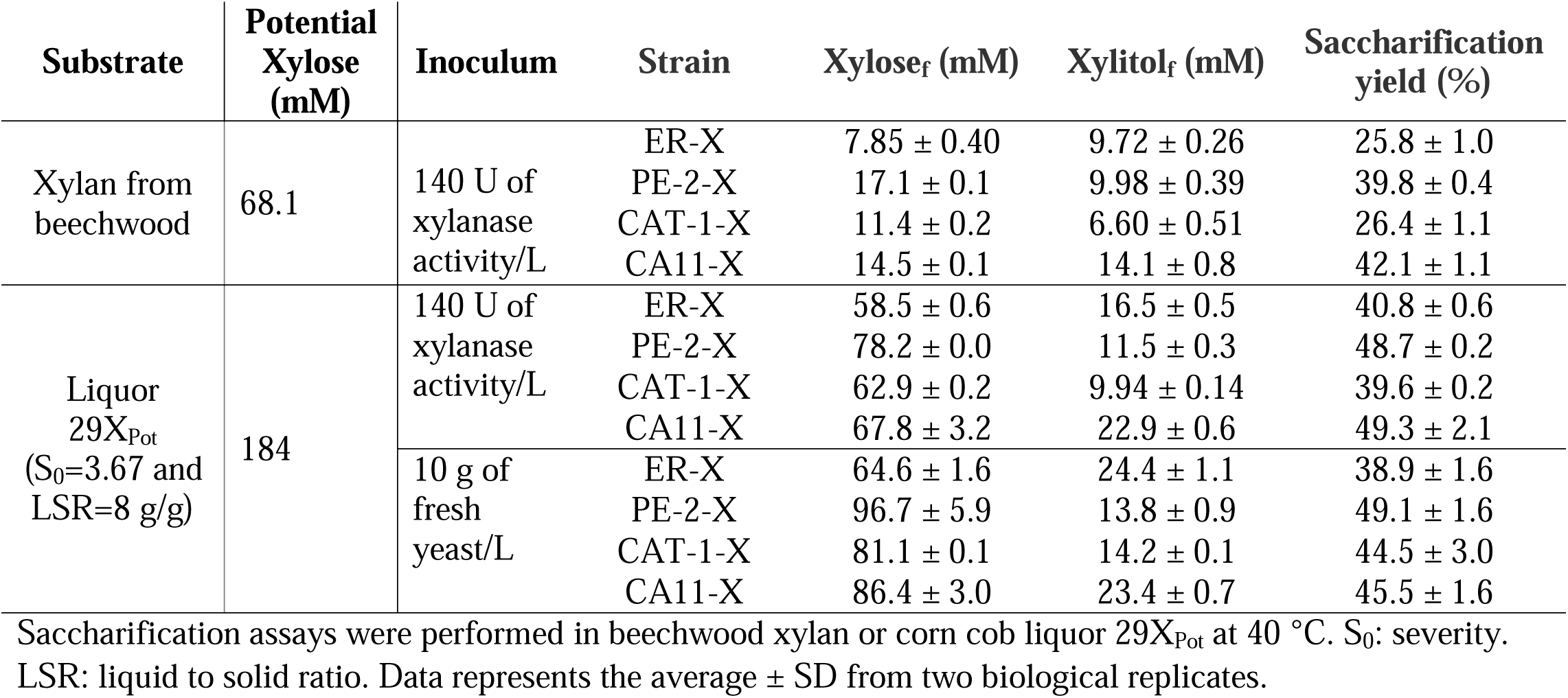
Saccharification parameters of the different *S. cerevisiae* strains displaying hemicellulolytic enzymes.

| Substrate | Potential Xylose (mM) | Inoculum | Strain | Xylose<sub>f</sub> (mM) | Xylitol<sub>f</sub> (mM) | Saccharification yield (%) |
| --- | --- | --- | --- | --- | --- | --- |
| Xylan from beechwood | 68.1 | 140 U of xylanase activity/L | ER-X | 7.85 ± 0.40 | 9.72 ± 0.26 | 25.8 ± 1.0 |
|  |  |  | PE-2-X | 17.1 ± 0.1 | 9.98 ± 0.39 | 39.8 ± 0.4 |
|  |  |  | CAT-1-X | 11.4 ± 0.2 | 6.60 ± 0.51 | 26.4 ± 1.1 |
|  |  |  | CA11-X | 14.5 ± 0.1 | 14.1 ± 0.8 | 42.1 ± 1.1 |
| Liquor 29X<sub>Pot</sub> (S<sub>0</sub>=3.67 and LSR=8 g/g) | 184 | 140 U of xylanase activity/L | ER-X | 58.5 ± 0.6 | 16.5 ± 0.5 | 40.8 ± 0.6 |
|  |  |  | PE-2-X | 78.2 ± 0.0 | 11.5 ± 0.3 | 48.7 ± 0.2 |
|  |  |  | CAT-1-X | 62.9 ± 0.2 | 9.94 ± 0.14 | 39.6 ± 0.2 |
|  |  |  | CA11-X | 67.8 ± 3.2 | 22.9 ± 0.6 | 49.3 ± 2.1 |
|  |  | 10 g of fresh yeast/L | ER-X | 64.6 ± 1.6 | 24.4 ± 1.1 | 38.9 ± 1.6 |
|  |  |  | PE-2-X | 96.7 ± 5.9 | 13.8 ± 0.9 | 49.1 ± 1.6 |
|  |  |  | CAT-1-X | 81.1 ± 0.1 | 14.2 ± 0.1 | 44.5 ± 3.0 |
|  |  |  | CA11-X | 86.4 ± 3.0 | 23.4 ± 0.7 | 45.5 ± 1.6 |

Saccharification assays were performed in beechwood xylan or corn cob liquor 29X<sub>Pot</sub> at 40 °C. S<sub>0</sub>: severity. LSR: liquid to solid ratio. Data represents the average ± SD from two biological replicates.

**Fig. 1.**
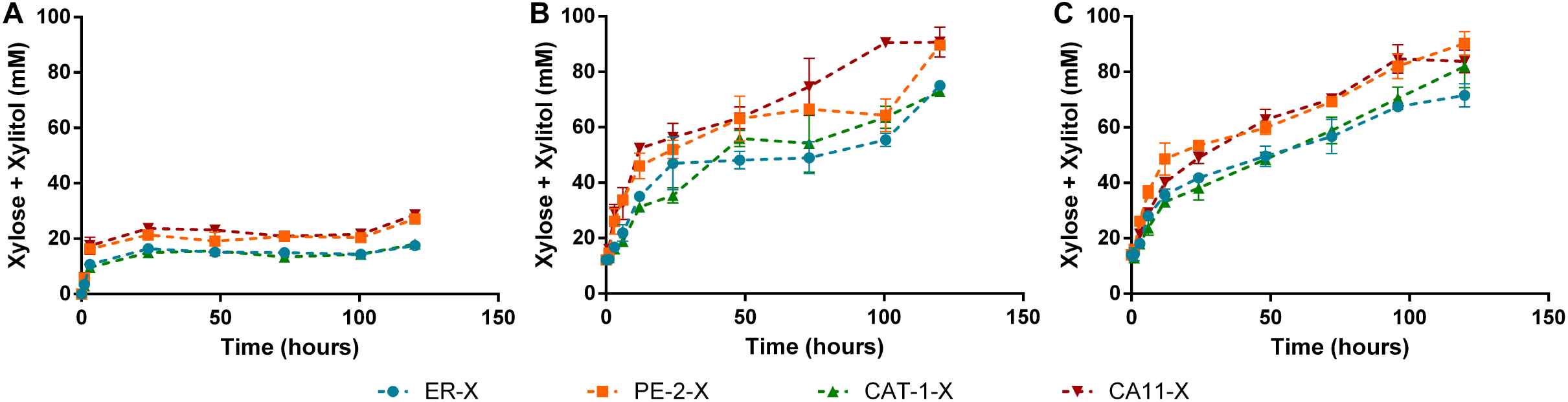
Saccharification capacity of the strains ER-X, PE-2-X, CAT-1-X and CA11-X. Assays were performed in xylan from beechwood (A) and in corn cob liquor 29X_Pot_ (B and C) at 40 °C. Inoculum for hydrolysis analyses was normalized by Units of xylanase activity (140 U/L; A and B) or by grams of fresh yeast (10 g/L; C). Data represents the average ± SD from at least two biological replicates.

As expected [21], the xylanase activity of the strains was higher at 40 °C than at 30 °C, being almost 2-fold higher for the ER-X strain (Table S1 [see Additional file 1]). Also, the ER-X strain presented the higher xylanase activity (Table S1 [see Additional file 1]). However, when evaluating the kinetic profile of saccharification of xylan from beechwood and XOS of corn cob liquor, PE-2-X and CA11-X were the strains capable of higher xylose release (measured as sum of xylose and xylitol, expressed in mM)(Fig. 1A, B and C). It should be noted that xylitol is produced from xylose by the action of unspecific aldose reductases natively present on *S. cerevisiae* strains and that this production varies among yeast strains [5, 16]. Thus, for analyzing xylan degrading capacity among different strains, the concentrations of xylose together with xylitol was considered (Fig. 1B, C and D). As seen, all the constructed strains were able to saccharify both substrates. Nevertheless, differences were observed among biocatalysts and substrates. The maximum saccharification of corn cob liquor was obtained with PE-2-X and CA11-X, reaching 49 % of the potential xylose, while ER-X reached only 39 % (Table 2). The incomplete hydrolysis can be a result of the heterogeneous structure of the beechwood/corn cob-derived xylan [22] in addition to product inhibition that may affect the cell-surface displayed enzymes due to xylose accumulation [7]. The hydrolysis of xylan is also known to be dependent of the raw material of origin and extraction method, varying in molecule length, degree of branching and presence of side groups [23]. This can be clearly observed in the different saccharification profiles observed between the commercial beechwood-derived xylan and the corn cob liquor (Fig. 1), with the maximum saccharification of beechwood xylan being only 42 % (Table 2). In fact, the total degradation of xylan requires the action of xylanolytic enzymes other than xylanase and xylosidade to remove side chains (such as arabinose, glucuronic acid, acetyl groups) from the xylan backbone. Nevertheless, the activity of these accessory enzymes, while increasing the amount of released xylose, would also cleave acetyl groups which would result in acetic acid accumulation in the medium with the consequent negative effects on yeast viability [20]. Different inoculum of the strains were also tested in corn cob liquor 29X_Pot_: in one experiment the strains were inoculated by xylanase activity (Fig. 1B) and in the other by wet cell weight (Fig. 1C). The similar xylan-degrading abilities observed with the same strain in different inoculums concentrations, also indicates that there is no enzyme/substrate ratio limitation.

Differences between the hydrolytic performance of the different strains (Fig. 1) were expected, as the efficiency of cell-surface display is dependent of several host-related factors [24], which have already been reported to vary among *S. cerevisiae* strains: *e*.*g*. secretory capacity [18], cell wall composition [25, 26], expression levels of host-cell genes [27]. Furthermore, yeast cell size is another source of variability, as larger cells have a lower total superficial cell area available for enzyme display per unit biomass. It should also be noted that the strains presenting higher hemicellulolytic capacity (CA11-X and PE-2-X, Fig. 1) were not the ones presenting higher XYLA and XYN activity values (Table S1 [see Additional file 1]), supporting the necessity of making the selection of appropriate yeast strain backgrounds in process-like conditions. Another factor to be taken into consideration is the flocculation ability, a process that is mediated by lectin-like receptors present in the cell surface which bind mannose residues in adjacent cells creating clusters of thousands of cells [28], but have also been reported to bind to a wide range of sugars [25]. Accordingly, the CA11 strain, being a flocculant strain, will present an high number of lectin-like receptors in the cell surface [25], that may potentiate the binding of the yeast to sugar residues in xylan/XOS, and benefit the hydrolysis in the long term due to substrate proximity. Furthermore, different predisposition to convert xylose into xylitol, normally observed between *S. cerevisiae* strains [16], may alleviate product inhibition of xylanase and xylosidase by xylose, increasing the overall degradation of xylan.

### Evaluation of engineered yeast strains’ capacity for direct production of ethanol from hemicellulosic liquor

To further evaluate the potential of the different yeast strains to directly produce ethanol from hemicellulose, *i*.*e*. degrade xylan and ferment the resulting xylose, they must be modified to be able to consume xylose. Recently, the oxidoreductase (xylose reductase-XR/xylitol dehydrogenase-XDH [29]) and the isomerase (xylose isomerase-XI [30]) xylose-consumption pathways were simultaneously expressed in robust industrial *S. cerevisiae* strains, improving the ethanol yield from a non-detoxified corn cob hydrolysate, with coupled higher furan detoxification and lower xylitol production, in comparison with the single expression of XR/XDH or XI [31]. Thus, in this work we opt for the simultaneous expression of the oxidoreductase and the isomerase metabolic pathways. Furthermore, to favour both saccharification and fermentation, the strains should be metabolic active at the optimal conditions previously described for the cell-surface displayed XYN, namely pH 5 and temperature of 40 °C [21]. In fact, the ability to ferment at higher temperatures has already been identified as a crucial trait for CBP yeast [1], and in this sense, the use of thermotolerant industrial *S. cerevisiae* strains is an advantage. Considering the trait variability among industrial strains, and the recognized necessity of a tailor-made design of yeast (considering process conditions and specific raw material) [16, 32], the strains ER-X, PE-2-X, CAT-1-X and CA11-X were evaluated in terms of capacity to ferment an hydrothermally pretreated corn cob liquor (supplemented with synthetic glucose) at 40 °C (Fig. S1 [see Additional file 1]). All the strains were capable of fermenting in these conditions, however the PE-2-X strain produced lower ethanol titers (Fig. S1[see Additional file 1]), being incapable of consuming all the glucose present in the medium (data not shown). Considering this poor performance, only the ER-X, CAT-X and CA11-X strains were further modified with the oxidoreductase and isomerase pathways for xylose consumption, resulting in strains ER-X-2P, CAT-X-2P and CA11-X-2P. Also important to note is the fact that the growth ability of the strains was not affected by the cell-surface display of the hemicellulases (Fig. S2 [see Additional file 1]) and that the integrations at the δ-sequences were found to be stable (Fig. S3 [see Additional file 1]).

The ER-X-2P, CAT-X-2P and CA11-X-2P strains were capable of producing ethanol directly from the corn cob liquor 29X_Pot_ at 40 °C (Fig. 2, Table 2), with ER-X-2P reaching the highest ethanol titer of 6.51 g/L (corresponding to an ethanol yield of 0.247 g per g of potential sugar), consuming almost all the xylose that was liberated in the medium (Fig. 2A, Table 2). On the other hand, the CAT-X-2P and CA11-X-2P strains produced significantly lower levels of ethanol (Fig. 2B and 2C, Table 2), being observable a clear accumulation of xylose at 24 hours of fermentation, result of the low fermentative aptitude of these strains in comparison with their xylan-degrading capacity. This superior performance of ER-X-2P is in accordance with previous reports of Ethanol Red excellent fermentation capacity, robustness, stress [33] and temperature tolerance [34]. In fact, Ethanol Red has recently been successively used as host for consolidated bioprocessing (CBP) for first generation bioethanol using raw starch [1], with the present work showing this strain applicability also for a more challenging CBP for second generation hemicellulosic ethanol.

**Fig. 2.**
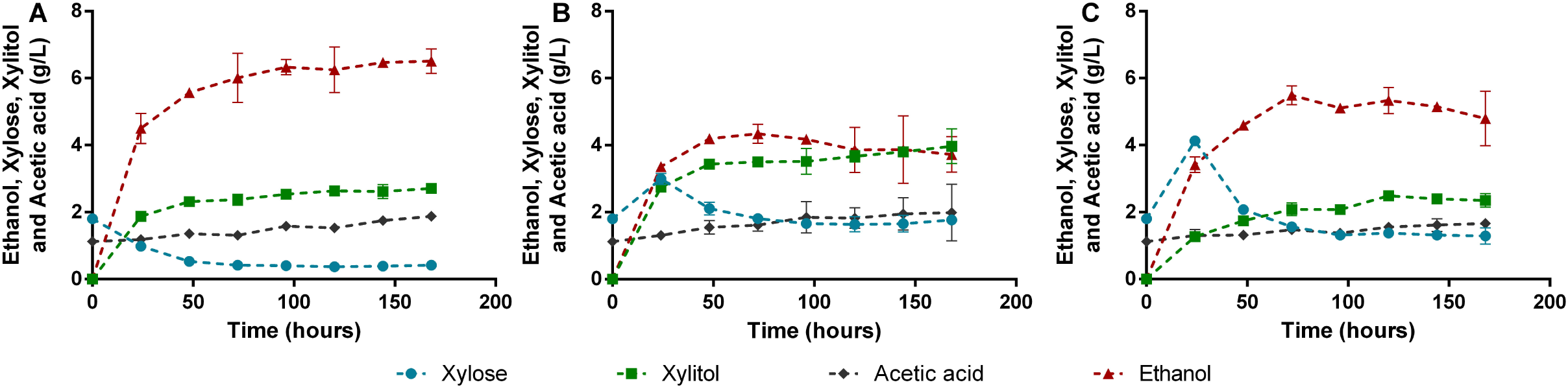
Consolidated bioprocessing profiles of the strains ER-X-2P (A), CAT-1-X-2P (B) and CA11-X-2P (C). Assays were performed in corn cob liquor 29X_Pot_ with an inoculum of 50 g/L fresh yeast. Data represents the average ± SD from two biological replicates.

### Consolidated bioprocessing of corn cob hemicellulosic liquor: evaluation of pretreatment conditions and inoculum size

As observed in Table 1, the increase of severity of treatment implies a higher solubilization of xylan in the liquid phase as XOS and xylose [35] which implies higher potential sugars to be fermented into ethanol. Therefore, the corn cob liquor obtained at severity of 3.99 (Liquor 32X_Pot_) was used to evaluate the role of inoculum size of ER-X-2P for simultaneous saccharification of XOS and fermentation into ethanol (Fig. 3A and B). In fact, this increase in pretreatment severity (3.99) allowed a higher ethanol production of 8.15 g/L, when compared with the 6.51 g/L obtained with the same strain and inoculum in liquor 29X_Pot_ (Fig. 3A and 2A, same strain and inoculum quantity). Furthermore, it was observed that increasing the inoculum from 50 to 100 g wet cells/L, an inoculum concentration in the order of magnitude of the normally reported for CBP strains [8, 36], improved the overall process (Fig. 3A and 3B) and increased the ethanol titer from 8.15 to 11.1 g/L (corresponding to an increase of 36 % in ethanol concentration) and the ethanol yield from 0.240 to 0.328 g per g of potential sugar (Table 3). These results represent the highest ethanol concentration obtained so far directly from an hemicellulosic liquor through a CBP microorganism. In fact, the ethanol titer attained with this CBP approach is superior to the ones previously reported for fermentation of non-detoxified corn cob hydrolysates obtained by acid hydrolysis [16, 31, 37]. Furthermore, the ethanol yield of 0.328 g/g is comparable to results previously obtained from simultaneous saccharfication and co-fermentation (SSCF) of a whole slurry (containing cellulose and hemicellulose and using commercial cellulases and hemicellulases) with an Ethanol Red strain modified for xylose consumption (yields of 0.32 and 0.28 g/g at 32 and 39 °C, respectively) [33].

**Table 3.**
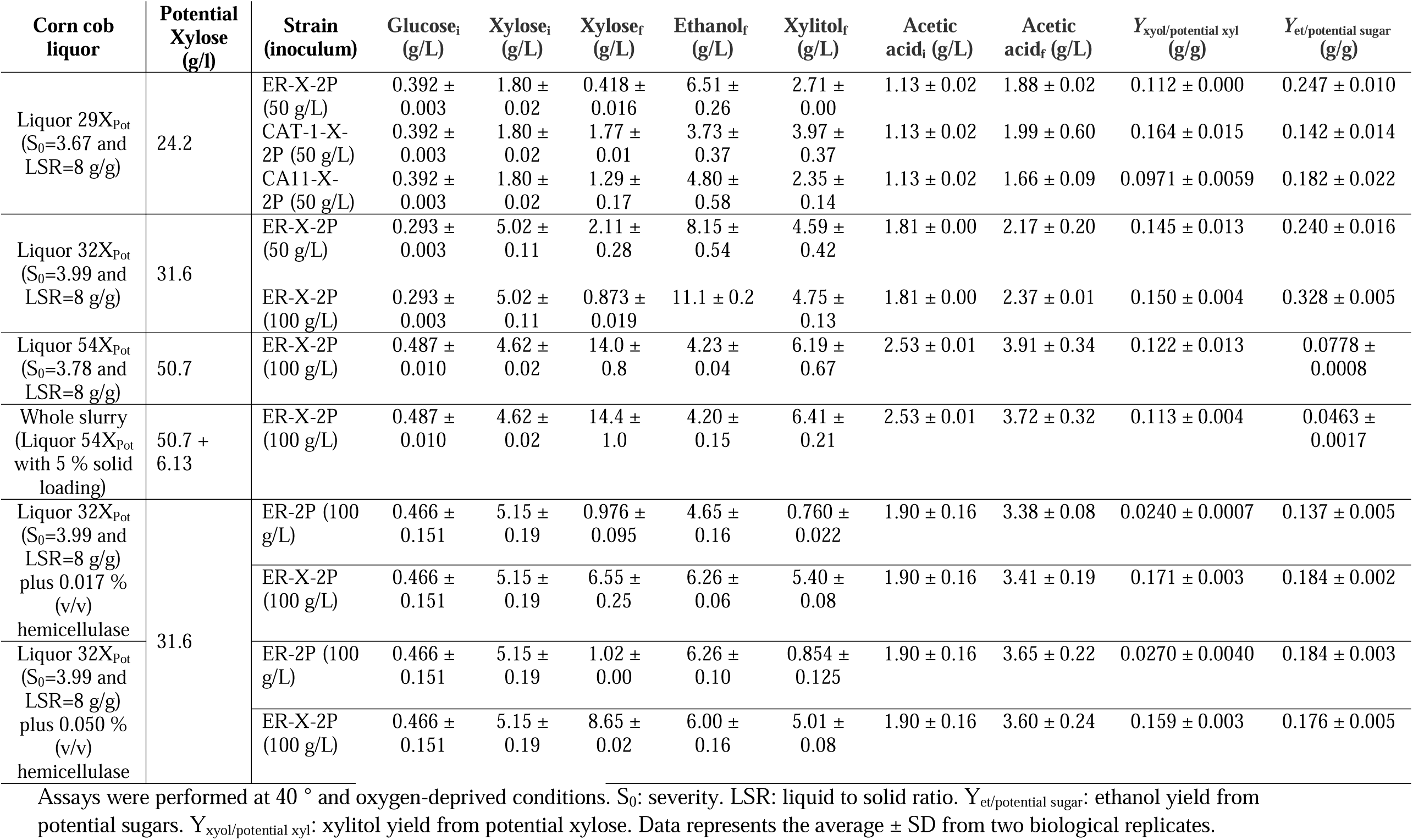
CBP and SSF parameters of the different xylose-consuming *S. cerevisiae* strains in corn cob liquors.

**Fig. 3.**
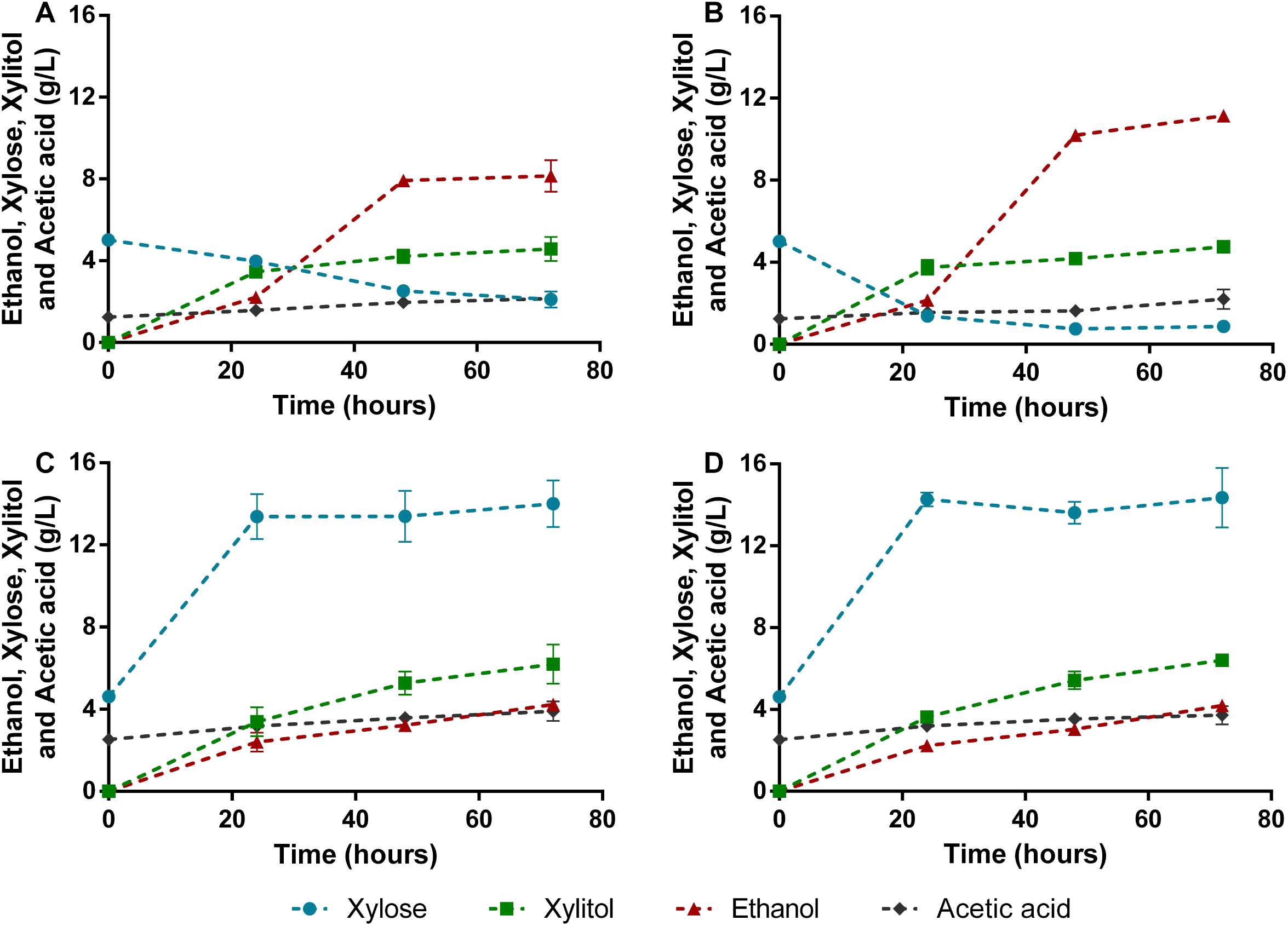
Consolidated bioprocessing profiles of the strain ER-X-2P. Assays were performed in corn cob liquor 32X_Pot_ (A and B), corn cob liquor 54X_Pot_ (C) and corn cob liquor 54X_Pot_ with 5 % solids (D) with an inoculum of 50 g/L fresh yeast (A) or 100 g/L fresh yeast (B, C and D). Data represents the average ± SD from two biological replicates.

Another strategy tested to increase ethanol concentration was to increase the potential of fermentable xylose derived from the corn cob pretreatment by reducing the liquid to solid ratio to 4 g/g, allowing a recovery of 48.8 g/L of XOS (Liquor 54X_Pot_, Table 1). However, the CBP of this liquor with ER-X-2P (Fig. 3C) resulted in lower ethanol titers than the ones previously obtained (even when using lower inoculums, Fig. 2A, 3A and 3B). This poor fermentative performance is explained by a significant increase in the inhibitory composition of the liquor resulting from the decrease in LSR, with liquor 54X_Pot_ presenting more than 2-fold the concentrations of acetic acid, HMF and furfural when comparing to liquor 29X_Pot_ (Table 1). Considering these, the pretreatment conditions for this process must be carefully defined in order to maximize xylan extraction while maintaining inhibitors concentration at a non-toxic level for the yeast strain.

Additionally, CBP with ER-X-2P was also performed by mixing corn cob liquor 54X_Pot_ with corn cob pretreated solids (obtaining a whole slurry as substrate), which contain 10.8 g of xylan per 100 g of pretreated corn cob (Table 1, Fig. 3D). Similar saccharification and fermentation profiles were observed for both experiments (with and without solids addition (Fig. 3C and D). This behavior shows that this strain was incapable of degrading pretreated corn cob solid. Nevertheless, there is no unproductive binding of the cell-surface-displayed enzymes to the lignin present in the solid fraction, a common problem in hydrolysis of lignocellulosic biomass [38], since equal saccharification yields from liquid phase (containing XOS) were obtained. With these results this strain is shown to be suitable for applications in integrated lignocellulosic processes (e.g. co-culture with cellulolytic strains for whole slurry CBP). Furthermore, this cell-surface display approach to CBP also facilitates the recycling of hydrolytic enzymes, an important technology for the development of economically viable processes [39]. In fact, cell recycling was attempted in CBP with ER-X-2P, and despite the gradual loss of yeast cell fermentative capacity, the hydrolytic activity of the cell-surface displayed enzymes was maintained after 2 cycles of recycling, releasing similar concentrations of xylose (Fig. S4 [see Additional file 1]). This also shows the advantages of the cell-surface display approach to express enzymes, and the suitability of these modified strains to function as whole-cell biocatalysts even with the loss of its native metabolic activity.

### Evaluation of commercial hemicellulase addition

The SSF of corn cob liquor 32X_Pot_ was also performed with the ER-2P strain (modified for xylose consumption with the XR/XDH and XI pathways but without the cell-surface display of enzymes) and with the addition of reduced quantities of the commercial cocktail of hemicellulases Cellic HTec2 (Fig. 4A and 4C), to compare with the direct production of ethanol from hemicellulose with only the ER-X-2P strain (Fig. 3B). Two concentrations of commercial hemicellulases were tested: 0.050 % (v/v), to equal the xylanase activity normally added with the inoculum of ER-X-2P; and 0.017 % (v/v), to evaluate the effect of a two third reduction of enzyme quantities (done to reduce process costs). Nevertheless, even with the higher concentration of commercial hemicellulase, the ethanol titers obtained with this method were lower than with the CBP process, 4.65 and 6.26 g/L compared to the previously attained 11.1 g/L (Table 3). Considering the high levels of xylobiose and xylotriose accumulated and the low levels of xylose throughout the experiments (Fig. 4A and 4C), it is clear that the commercial cocktail, while being capable of producing XOS with low degree of polymerization, is incapable to release xylose at the rate that it is being consumed by ER-2P. This supports the already reported deficit of xylosidase activity in commercial hemicellulase cocktails [8, 40]. On the other hand, when performing the SSF with addition of commercial cocktail but with the ER-X-2P strain (Fig. 4B and 4D), the concentration of xylobiose and xylotriose was maintained low throughout the experiment, showing that the xylanolytic strain compensates the low xylosidase activity of the commercial hemicellulases. Furthermore, the important role of ER-X-2P for the degradation of hemicellulose is also shown, as it produced similar ethanol titers in SSF with both concentrations of commercial hemicellulases (Table 3, Fig. 4B and 4D), while with ER-2P the ethanol titer was significantly lower when less hemicellulase cocktail was added (Table 3, Fig. 4A and 4C). Nevertheless, all the SSF experiments of corn cob liquor 32X_Pot_ with addition of commercial hemicellulases resulted in low ethanol titers (≤ 6.26 g/L, Fig. 4), while the CBP of the same liquor with only ER-X-2P produced at least 1.77-fold higher concentrations (Fig. 3B, Table 3). This low performance, may be attributed to the levels of acetic acid that were released throughout the experiment (mainly in the first 24 h, Fig. 4), due to the fact that, in contrary to the ER-X-2P strain, the commercial cocktail contains other hemicellulases capable of removing xylan side chains, such as acetyl xylan esterase (EC 3.1.1.72). In fact, while in the CBP of liquor 32X_Pot_ with no enzyme addition,the increase of this weak acid was minimal (1.3-fold, Fig. 3B), with commercial enzymes supplementation the acetic acid concentrations increased more than 1.8-fold in the 72 h of SSF (Fig. 4), which clearly had a negative impact in the yeast cell fermentative capacity. Summing up, the use of this commercial cocktail in conjugation with ER-X-2P strain, while increasing the amount of released xylose, also significantly increased the acetic acid concentration of the medium, reducing the overall ethanol production in comparison with CBP using only the yeast strain ER-X-2P (Fig. 5). In fact, an overall balance of corn cob processing for hemicellulosic ethanol production considering the results obtained in this work with Liquor 32X_Pot_ (Fig. 5) clearly shows the advantage of CBP: while the SSF process may produce a maximum of 57.8 kg of ethanol from 1 ton of corn cob requiring 1.9 kg of commercial enzymatic cocktail, the sole use of ER-X-2P yeast strain as biocatalyst allows the production of 102.8 kg of ethanol from the same amount of raw material, without addition of exogenous enzymes. Additionally, the SSF with the ER-2P strain allows the attainment of only 42.9 kg of ethanol/ton of corn cob, also highlighting the importance of using the hemicellulolytic strains. These results show that the use of engineered industrial *Saccharomyces cerevisiae* strains as whole-cell biocatalysts for the saccharification of corn cob-derived hemicellulosic liquor is advantageous for ethanol production in comparison with the use of commercial enzymatic cocktails.

**Fig. 4.**
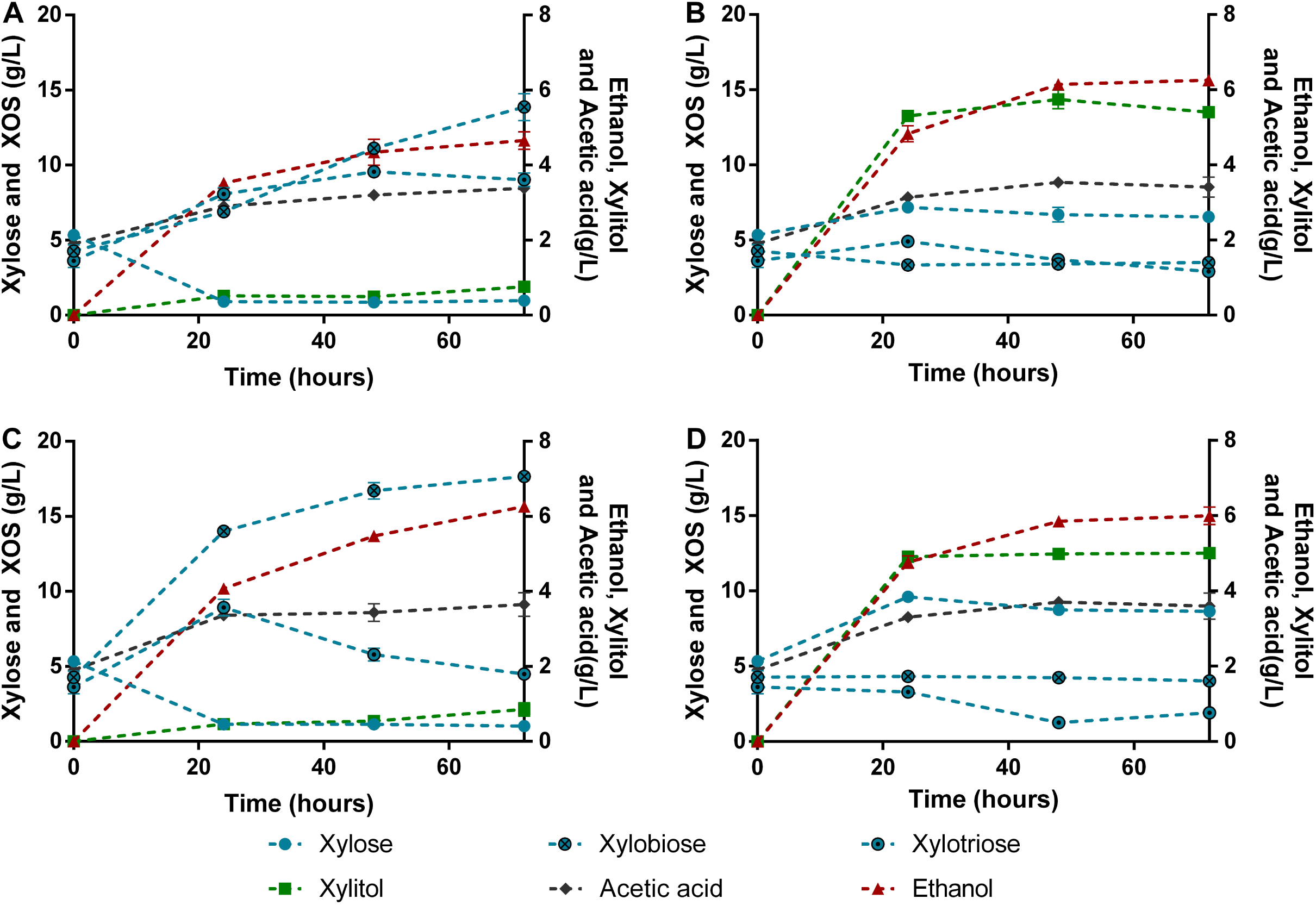
Simultaneous saccharification and fermentation profiles of the strains ER-2P (A and C) and ER-X-2P (B and D). Assays were performed in corn cob liquor 32X_Pot_ with addition of commercial hemicellulase (0.017 % (v/v): A and B; 0.050 % (v/v): C and D) with an inoculum of 100 g/L fresh yeast. Data represents the average ± SD from two biological replicates.

**Fig. 5.**
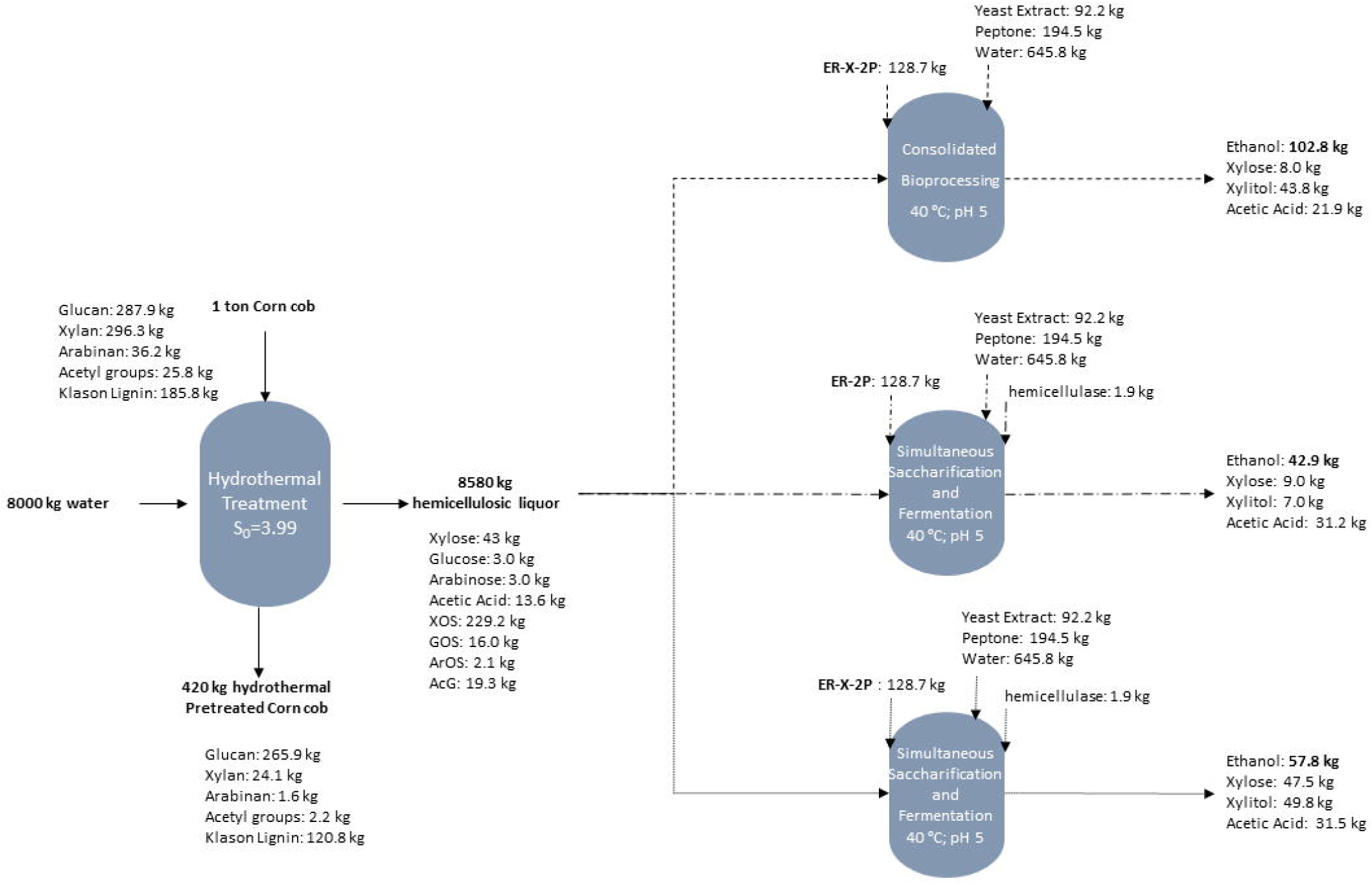
Mass balance of consolidated bioprocessing and simultaneous saccharification and fermentation of the hemicellulosic liquor 32X_Pot_.

## Conclusions

These results show the highest ethanol concentration reported from direct conversion of hemicellulosic liquors by *S. cerevisiae*, which are also higher than ethanol values obtained from corn cob-derived hemicellulose after acid post hydrolysis. In this sense, the use of engineered industrial *Saccharomyces cerevisiae* strains as whole-cell biocatalysts is shown to be a suitable alternative to commercial enzymatic cocktails or chemical hydrolysis and represents a greener and economical approach to produce hemicellulosic ethanol from corn cob biomass. Additionally, the potential of robust industrial *S. cerevisiae* strains as hosts for the design of whole-cell biocatalysts is demonstrated, paving the way for the construction of more efficient strains for consolidated bioprocessing in lignocellulosic biorefineries.

## Methods

### Yeast strains and plasmid construction

*Saccharomyces cerevisiae* strains and main plasmids used in this work are listed in Table 4 and primers and plasmids used for cloning steps are presented on Table S2 [see Additional file 1]. *Escherichia coli* DH5α was used for plasmid construction, propagation and maintenance. The integrative plasmid pI23-BGL1-kanMX for expression of *Aspergillus aculeatus* β-glucosidase 1 (BGL1; EC 3.2.1.21; Accession P48825.1) was obtained from the plasmid pIBG-SSA by substitution of the *HIS3* for the integration region 23 (3′ non-coding region of genes *YCL054W* and *YCL052C*, chromosome III) [41] and the *kanMX* resistance marker. The plasmid pI5-XylA-NatX for expression of *Aspergillus oryzae* β-xylosidase A (XYLA; EC 3.2.1.37; Accession OOO12801.1) was obtained from the plasmid pIK-BX-SSS by substitution of the *LYS2* for the integration region I5 (3′ non-coding region of genes *YLL055W* and *YLL054C*, chromosome XII) [3] and the *natMX* resistance marker. The δ-integration plasmid pδW-XYN-kanMX for expression of *Trichoderma reesei* endoxylanase II (XYN; EC 3.2.1.8; Accession XP_006968947.1) was constructed from the pδW-EX-SSS by substitution of the *TRP1* marker for the kanMX-UkG1 cassette, containing both the *kanMX* resistance marker and the green fluorescent protein mUkG1 [42] for monitoring copy number integration. Plasmid assembling steps were performed with the In-Fusion HD Cloning Kit (Clontech, Mountain View, CA, USA). The gene cassettes used for the cell-surface expression of hemicellulases were previously optimized [43, 44] and consist of: coding sequences of *SED1* promoter, *SED1* secretion signal, and *SAG1* anchoring domain for BGL1 expression; and coding sequences of *SED1* promoter, *SED1* secretion signal, and *SED1* anchoring domain for expression of XYLA and XYNII. The plasmids pI23-BGL1-kanMX, pI5-XylA-NatMX and pδW-XYN-kanMX were linearized with BstZ17I, SpeI and AscI, respectively, and were sequentially transformed by lithium acetate [45] and integrated in the different yeast strains. The transformants were selected on YPD plates (10 g/L yeast extract, 20 g/L peptone, 20 g/L glucose and 20 g/L agar) containing 300 μg/mL of G418 (pI23-BGL1-kanMX and pδW-XYN-kanMX) or 100 μg/mL of clonNAT (pI5-XylA-NatMX) and were preserved at 4 °C on YPD plates. Before integration of pδW-XYN-kanMX, the *kanMX* marker inserted with the pI23-BGL1-kanMX integration was removed using the CRE-loxp recombinase system [46]. After each integration, transformants of each of the different yeast hosts were screened, and the transformant presenting higher enzymatic activity was selected for further work (data not shown), and were denominated ER-X, PE-2-X, CAT-1-X and CA11-X. For xylose consumption the plasmid pMEC1049+XI, for expression of both the oxidoreductase and the isomerase pathways [31], was introduced by lithium acetate transformation [45] on the yeast strains displaying the hemicellulolytic enzymes, originating the strains ER-X-2P, CAT-1-X-2P and CA11-X-2P, and also on the ER wild type strain, originating the strain ER-2P. The transformants were selected on YPD plates containing 300 μg/mL of hygromycin and were preserved at 4 °C on YPX plates (10 g/L yeast extract, 20 g/L peptone, 20 g/L xylose and 20 g/L agar) containing 300 μg/mL of hygromycin to maintain selective pressure.

**Table 4.**
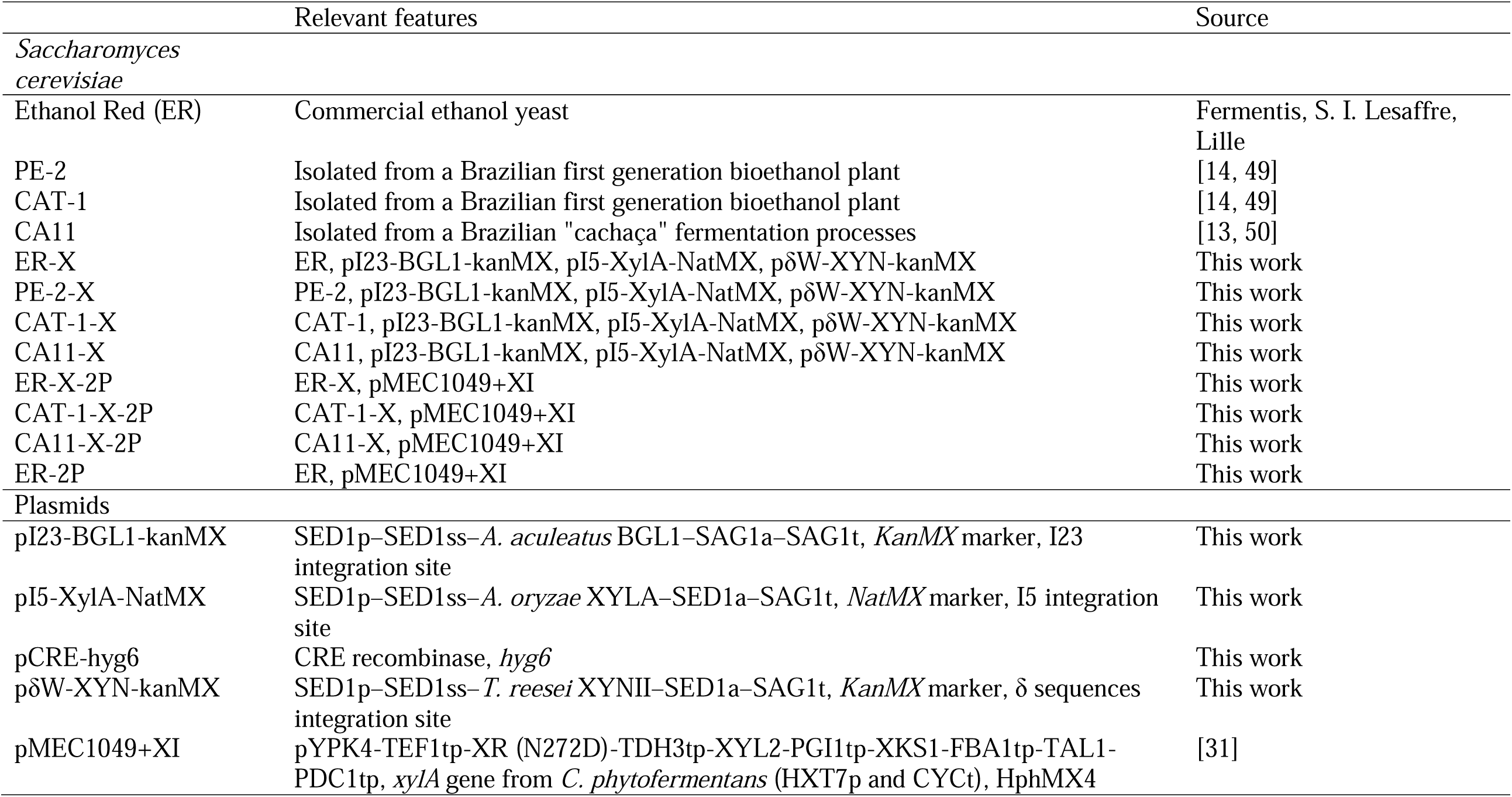
*Saccharomyces cerevisiae* strains and main plasmids used in this study.

|  | Relevant features | Source |
| --- | --- | --- |
| *Saccharomyces cerevisiae* |  |  |
| Ethanol Red (ER) | Commercial ethanol yeast | Fermentis, S. I. Lesaffre, Lille |
| PE-2 | Isolated from a Brazilian first generation bioethanol plant | [14, 49] |
| CAT-1 | Isolated from a Brazilian first generation bioethanol plant | [14, 49] |
| CA11 | Isolated from a Brazilian "cachaça" fermentation processes | [13, 50] |
| ER-X | ER, pI23-BGL1-kanMX, pI5-XylA-NatMX, pδW-XYN-kanMX | This work |
| PE-2-X | PE-2, pI23-BGL1-kanMX, pI5-XylA-NatMX, pδW-XYN-kanMX | This work |
| CAT-1-X | CAT-1, pI23-BGL1-kanMX, pI5-XylA-NatMX, pδW-XYN-kanMX | This work |
| CA11-X | CA11, pI23-BGL1-kanMX, pI5-XylA-NatMX, pδW-XYN-kanMX | This work |
| ER-X-2P | ER-X, pMEC1049+XI | This work |
| CAT-1-X-2P | CAT-1-X, pMEC1049+XI | This work |
| CA11-X-2P | CA11-X, pMEC1049+XI | This work |
| ER-2P | ER, pMEC1049+XI | This work |
| Plasmids |  |  |
| pI23-BGL1-kanMX | SED1p–SED1ss–*A. aculeatus* BGL1–SAG1a–SAG1t, *KanMX* marker, I23 integration site | This work |
| pI5-XylA-NatMX | SED1p–SED1ss–*A. oryzae* XYLA–SED1a–SAG1t, *NatMX* marker, I5 integration site | This work |
| pCRE-hyg6 | CRE recombinase, *hyg6* | This work |
| pδW-XYN-kanMX | SED1p–SED1ss–*T. reesei* XYNII–SED1a–SAG1t, *KanMX* marker, δ sequences integration site | This work |
| pMEC1049+XI | pYPK4-TEF1tp-XR (N272D)-TDH3tp-XYL2-PGI1tp-XKS1-FBA1tp-TAL1-PDC1tp, *xylA* gene from *C. phytofermentans* (HXT7p and CYCt), HphMX4 | [31] |

### Enzymatic activities

For determination of enzymatic activities the yeast cells were grown in YPD medium (10 g/L yeast extract, 20 g/L peptone and 20 g/L glucose) for 48 h at 30 °C (Table S1 [see Additional file 1]) or YPX medium (10 g/L yeast extract, 20 g/L peptone and 20 g/L xylose) for 48 h at 35 °C for xylanase activity determinations of the inoculums for SSF experiments (data not shown); the cells were collected by centrifugation at 1000 g for 5 min. β-glucosidase 1 and β-xylosidase A activities were measured at 30 °C with nitrophenyl-β-d-glucopyranoside and p-nitrophenyl-β-D-xylopyranoside (Nacalai Tesque, Inc., Kyoto, Japan), respectively, as previously described [47]. For the measurement of xylanase activity, the cells (20 g wet cells/L) were incubated in 10 g/L of xylan from beechwood (Sigma, ≥ 90 % purity) in 50 mM sodium acetate buffer (pH 5.0) for 10 min at 250 rpm orbital agitation at 30 and 40 °C. The amount of reducing sugar released from the substrate was measured by the DNS method [48]. One unit of xylanase activity was defined as the amount of enzyme required to release 1 μmol of reducing sugar per minute. Dry cell weight (DCW) of the yeast strains was estimated to be 0.15-fold (ER, PE-2 and CAT-1) and 0.18-fold (CA11) that of the wet cell weight.

### Preparation of corn cob liquor

Corn cob was collected, milled and submitted to hydrothermal treatment under non-isothermal conditions, selected based on previous works [5, 19], in a 2 L stainless steel reactor (Parr Instruments Company) equipped with Parr PDI temperature controller (model 4848) at liquid to solid ratio of 8 g distilled water/1 g of oven dry corn cob or of 4 g distilled water/1 g of oven dry corn cob (Table 3). T_max_ of 205 °C, 211 °C and 207 °C corresponding to a severity (S_0_) of 3.67, 3.99 and 3.78, were used to obtain the hemicellulosic liquors, that were denominated according to their potential in xylose (g/L), i.e. Liquor29X_Pot_, 32X_Pot_ and 54X_Pot_, respectively. After treatment, the resulting solid and liquid (liquor) phases were separated by filtration. Solid phase was recovered and washed for solid yield determination. Severity (S_0_) was calculated by the following equation:

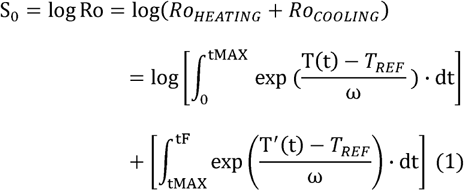

where R_0_ is the severity factor, t_MAX_ (min) is the time needed to achieve the target temperature T_MAX_ (°C), t_F_ (min) is the time needed for the whole heating–cooling period, and T(t) and T’(t) represent the temperature profiles in the heating and cooling stages, respectively. Calculations were made using the values reported usually for ω and T_REF_ (14.75 °C and 100 °C, respectively).

### Saccharification of xylan and xylooligosaccharides from corncob liquors

Yeast cells used for saccharification were cultivated at 30 °C for 48 h, with orbital agitation (200 rpm), in YPD medium. Cells were washed with water and the saccharification assays were inoculated with 10 g of wet yeast/L or with the biomass corresponding to 140 Units of xylanase activity/L. Saccharification was carried out in 50 mL Erlenmeyer flasks, with cotton stopper and working volume of 15 mL, in 10 g/L of xylan from beechwood in 50 mM sodium acetate buffer (pH 5.0) or in corn cob liquor 29X_Pot_ (pH adjusted to 5.0). Assays were performed at 40 °C in an orbital shaker at 150 rpm and samples were collected throughout the experiment for high-performance liquid chromatography (HPLC) analysis.

### Consolidated bioprocessing (CBP) and Simultaneous Saccharification and Fermentation (SSF) assays

Yeast cells used for CBP/SSF were cultivated at 35 °C for 48 h, with orbital agitation (200 rpm), in YPX medium. Cells were washed with NaCl (0.9 %) and the CBP/SSF assays were inoculated with 50 or 100 g of wet yeast/L. CBP/SSF was carried out in 100 mL Erlenmeyer flasks, with glycerol lock to create oxygen-deprived conditions and working volume of 30 mL. Media consisted of corn cob liquor 29X_Pot_, 32X_Pot_ or 54X_Pot_; or corn cob liquor 54X_Pot_ with 5 % (DCW/v) of pretreated corn cob solid fraction. All media were supplemented with YP (10 g/L yeast extract, 20 g/L peptone) and adjusted to pH 5.0. SSF of corn cob liquor 32X_Pot_ was performed with addition of 0.017 % (v/v) or 0.050 % (v/v) of the commercial hemicellulase Cellic HTec2 (Novozymes). Assays were performed at 40 °C in an orbital shaker at 150 rpm and samples were collected throughout the experiment for HPLC analysis.

### Analytical methods

The chemical composition of raw material and solid phases obtained after pretreatment were determined following standard methods described by NREL protocols (NREL/TP-510-42618-42622-4218). The liquid phase was analyzed directly and after posthydrolysis (4 % H_2_SO_4_ at 121 °C for 20 min) allowing the quantification of monosaccharides, oligosaccharides (measured as monomers equivalents), acetic acid and furan derived compounds (furfural and HMF). Samples from corn cob treatment and from saccharification and fermentation assays were analyzed for quantification of glucose, xylose, xylitol, xylobiose, xylotriose, acetic acid, ethanol, HMF and furfural by HPLC using a Bio-Rad Aminex HPX-87H column, operating at 60 °C, with 0.005 M H_2_SO_4_ and at a flow rate of 0.6 mL/min. HMF and furfural were detected using an UV detector set at 270 nm, whereas the other compounds were detected using a Knauer-IR refractive index detector.

### Determination of fermentation parameters

Saccharification yield (%) was calculated using the following equation:

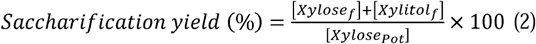

where [Xylose_f_] is xylose concentration (mM) at the end of saccharification assay, [Xylitol_f_] is xylitol concentration (mM) at the end of saccharification assay and [Xylose_Pot_] is the potential xylose (mM) present in the media used.

Ethanol yield from potential sugars (Y_et/potential sugar_) was calculated using the following equation:

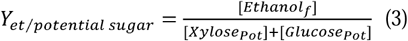

where [Ethanol_f_] is ethanol concentration (g/L) at the end of fermentation assay, [Xylose_Pot_] and [Glucose_Pot_] are the potential xylose (g/L) and glucose (g/L), respectively, present in the media used.

Xylitol yield from potential xylose (Y_xyol/potential xyl_) was calculated using the following equation:

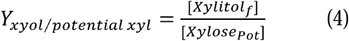

where [Xylitol_f_] is xylitol concentration (g/L) at the end of fermentation assay and [Xylose_Pot_] is the potential xylose (g/L) present in the media used.

## Acknowledgements

This work has been carried out at the Biomass and Bioenergy Research Infrastructure (BBRI)-LISBOA-01-0145-FEDER-022059, supported by Operational Programme for Competitiveness and Internationalization (PORTUGAL2020), by Lisbon Portugal Regional Operational Programme (Lisboa 2020) and by North Portugal Regional Operational Programme (Norte 2020) under the Portugal 2020 Partnership Agreement, through the European Regional Development Fund (ERDF) and has been supported by the Portuguese Foundation for Science and Technology (FCT) under the scope of the strategic funding of UIDB/04469/2020, the MIT-Portugal Program (Ph.D. Grant PD/BD/128247/2016 to Joana T. Cunha) and through Project FatVal (POCI-01-0145-FEDER-032506) and BioTecNorte operation (NORTE-01-0145-FEDER-000004) funded by the European Regional Development Fund under the scope of Norte2020 – Programa Operacional Regional do Norte.

## Additional file 1

Fermentation of ER-X, PE-2-X, CAT-1-X and CA11-X in in corn cob liquor supplemented with 100 g/L glucose at 40 °C; Growth profiles of ER-X, CAT-1-X and CA11-X and corresponding type strains in YPD; Stability evaluation of the integration at the δ-sequences; Fermentation of corn cob liquor with ER-X-2P strain and 2 cycles of cell recycling; Enzymatic activities; Plasmids and primers used for subcloning steps.

## Notes

### Competing Interest Statement

The authors have declared no competing interest.

## References

1. Cripwell RA, Rose SH, Favaro L, van Zyl WH. Construction of industrial *Saccharomyces cerevisiae* strains for the efficient consolidated bioprocessing of raw starch. Biotechnol Biofuels. 2019;12:201.

2. Loaces I, Schein S, Noya F. Ethanol production by *Escherichia coli* from *Arundo donax* biomass under SSF, SHF or CBP process configurations and in situ production of a multifunctional glucanase and xylanase. Bioresour Technol. 2017;224:307–313.

3. Liu Z, Inokuma K, Ho S-H, Haan Rd, Hasunuma T, van Zyl WH, et al. Combined cell-surface display- and secretion-based strategies for production of cellulosic ethanol with *Saccharomyces cerevisiae*. Biotechnol Biofuels. 2015;8:162–162.

4. Lin B, Tao Y. Whole-cell biocatalysts by design. Microb Cell Fact. 2017;16:106.

5. Baptista SL, Cunha JT, Romaní A, Domingues L. Xylitol production from lignocellulosic whole slurry corn cob by engineered industrial *Saccharomyces cerevisiae* PE-2. Bioresour Technol. 2018;267:481–491.

6. Hasunuma T, Hori Y, Sakamoto T, Ochiai M, Hatanaka H, Kondo A. Development of a GIN11/FRT-based multiple-gene integration technique affording inhibitor-tolerant, hemicellulolytic, xylose-utilizing abilities to industrial *Saccharomyces cerevisiae* strains for ethanol production from undetoxified lignocellulosic hemicelluloses. Microb Cell Fact. 2014;13:145.

7. Rohman A, van Oosterwijk N, Puspaningsih NNT, Dijkstra BW. Structural basis of product inhibition by arabinose and xylose of the thermostable GH43 β-1,4-xylosidase from *Geobacillus thermoleovorans* IT-08. PLoS One. 2018;13:e0196358.

8. Sakamoto T, Hasunuma T, Hori Y, Yamada R, Kondo A. Direct ethanol production from hemicellulosic materials of rice straw by use of an engineered yeast strain codisplaying three types of hemicellulolytic enzymes on the surface of xylose-utilizing *Saccharomyces cerevisiae* cells. J Biotechnol. 2012;158:203–210.

9. Katahira S, Fujita Y, Mizuike A, Fukuda H, Kondo A. Construction of a xylan-fermenting yeast strain through codisplay of xylanolytic enzymes on the surface of xylose-utilizing *Saccharomyces cerevisiae* cells. Appl Environ Microbiol. 2004;70:5407–5414.

10. Sun J, Wen F, Si T, Xu J-H, Zhao H. Direct conversion of xylan to ethanol by recombinant *Saccharomyces cerevisiae* strains displaying an engineered minihemicellulosome. Appl Environ Microbiol. 2012;78:3837–3845.

11. Lee S-M, Jellison T, Alper HS. Xylan catabolism is improved by blending bioprospecting and metabolic pathway engineering in *Saccharomyces cerevisiae*. Biotechnol J. 2015;10:575–575.

12. Sasaki Y, Takagi T, Motone K, Kuroda K, Ueda M. Enhanced direct ethanol production by cofactor optimization of cell surface-displayed xylose isomerase in yeast. Biotechnol Prog. 2017;33:1068–1076.

13. Pereira FB, Guimarães PMR, Teixeira JA, Domingues L. Selection of *Saccharomyces cerevisiae* strains for efficient very high gravity bio-ethanol fermentation processes. Biotechnol Lett. 2010;32:1655–1661.

14. Pereira FB, Guimarães PMR, Teixeira JA, Domingues L. Robust industrial *Saccharomyces cerevisiae* strains for very high gravity bio-ethanol fermentations. J Biosci Bioeng. 2011;112:130–136.

15. Pereira FB, Romaní A, Ruiz HA, Teixeira JA, Domingues L. Industrial robust yeast isolates with great potential for fermentation of lignocellulosic biomass. Bioresour Technol. 2014;161:192–199.

16. Costa CE, Romaní A, Cunha JT, Johansson B, Domingues L. Integrated approach for selecting efficient *Saccharomyces cerevisiae* for industrial lignocellulosic fermentations: Importance of yeast chassis linked to process conditions. Bioresour Technol. 2017;227:24–34.

17. Davison SA, den Haan R, van Zyl WH. Heterologous expression of cellulase genes in natural *Saccharomyces cerevisiae* strains. Appl Microbiol Biotechnol. 2016;100:8241–8254.

18. Davison SA, den Haan R, van Zyl WH. Identification of superior cellulase secretion phenotypes in haploids derived from natural *Saccharomyces cerevisiae* isolates. FEMS Yeast Res. 2019;19:

19. Rivas B, Dominguez JM, Dominguez H, Parajó JC. Bioconversion of posthydrolysed autohydrolysis liquors: an alternative for xylitol production from corn cobs. Enzyme Microb Technol. 2002;31:431–438.

20. Cunha JT, Romaní A, Costa CE, Sá-Correia I, Domingues L. Molecular and physiological basis of *Saccharomyces cerevisiae* tolerance to adverse lignocellulose-based process conditions. Appl Microbiol Biotechnol. 2019;103:159–175.

21. Fujita Y, Katahira S, Ueda M, Tanaka A, Okada H, Morikawa Y, et al. Construction of whole-cell biocatalyst for xylan degradation through cell-surface xylanase display in *Saccharomyces cerevisiae*. J Mol Catal B: Enzym. 2002;17:189–195.

22. Javed U, Aman A, Qader SAU. Utilization of corncob xylan as a sole carbon source for the biosynthesis of endo-1,4-β xylanase from *Aspergillus niger* KIBGE-IB36. Bioresour Bioprocessing. 2017;4:19.

23. Gírio FM, Fonseca C, Carvalheiro F, Duarte LC, Marques S, Bogel-Lukasik R. Hemicelluloses for fuel ethanol: A review. Bioresour Technol. 2010;101:4775–4800.

24. Tanaka T, Kondo A. Cell surface engineering of industrial microorganisms for biorefining applications. Biotechnol Adv. 2015;33:1403–11.

25. Nayyar A, Walker G, Wardrop F, Adya AK. Flocculation in industrial strains of *Saccharomyces cerevisiae*: role of cell wall polysaccharides and lectin-like receptors. J I Brewing. 2017;123:211–218.

26. Bou Zeidan M, Zara G, Viti C, Decorosi F, Mannazzu I, Budroni M, et al. L-histidine inhibits biofilm formation and FLO11-associated phenotypes in *Saccharomyces cerevisiae* flor yeasts. PLoS One. 2014;9:e112141.

27. Ibanez C, Perez-Torrado R, Morard M, Toft C, Barrio E, Querol A. RNAseq-based transcriptome comparison of *Saccharomyces cerevisiae* strains isolated from diverse fermentative environments. Int J Food Microbiol. 2017;257:262–270.

28. Masy CL, Henquinet A, Mestdagh MM. Fluorescence study of lectinlike receptors involved in the flocculation of the yeast *Saccharomyces cerevisiae*. Can J Microbiol. 1992;38:405–409.

29. Romani A, Pereira F, Johansson B, Domingues L. Metabolic engineering of *Saccharomyces cerevisiae* ethanol strains PE-2 and CAT-1 for efficient lignocellulosic fermentation. Bioresour Technol. 2015;179:150–158.

30. Brat D, Boles E, Wiedemann B. Functional expression of a bacterial xylose isomerase in *Saccharomyces cerevisiae*. Appl Environ Microbiol. 2009;75:2304–2311.

31. Cunha JT, Soares PO, Romaní A, Thevelein JM, Domingues L. Xylose fermentation efficiency of industrial *Saccharomyces cerevisiae* yeast with separate or combined xylose reductase/xylitol dehydrogenase and xylose isomerase pathways. Biotechnol Biofuels. 2019;12:20.

32. Cunha JT, Aguiar TQ, Romani A, Oliveira C, Domingues L. Contribution of *PRS3, RPB4* and *ZWF1* to the resistance of industrial *Saccharomyces cerevisiae* CCUG53310 and PE-2 strains to lignocellulosic hydrolysate-derived inhibitors. Bioresour Technol. 2015;191:7–16.

33. Demeke MM, Dietz H, Li Y, Foulquié-Moreno MR, Mutturi S, Deprez S, et al. Development of a D-xylose fermenting and inhibitor tolerant industrial *Saccharomyces cerevisiae* strain with high performance in lignocellulose hydrolysates using metabolic and evolutionary engineering. Biotechnol Biofuels. 2013;6:89.

34. Pinheiro T, Lip KYF, García-Ríos E, Querol A, Teixeira J, van Gulik W, et al. Differential proteomic analysis by SWATH-MS unravels the most dominant mechanisms underlying yeast adaptation to non-optimal temperatures under anaerobic conditions. bioRxiv. 2020;

35. Garrote G, Domi, x, nguez H, Parajó JC. Autohydrolysis of corncob: study of non-isothermal operation for xylooligosaccharide production. Journal of Food Engineering. 2002;52:211–218.

36. Liu Z, Inokuma K, Ho S-H, den Haan R, van Zyl WH, Hasunuma T, et al. Improvement of ethanol production from crystalline cellulose via optimizing cellulase ratios in cellulolytic *Saccharomyces cerevisiae*. Biotechnol Bioeng. 2017;114:1201–1207.

37. Lee J-W, Zhu JY, Scordia D, Jeffries TW. Evaluation of ethanol production from corncob using *Scheffersomyces (Pichia) stipitis* CBS 6054 by volumetric scale-up. Appl Biochem Biotechnol. 2011;165:814–822.

38. Saini JK, Patel AK, Adsul M, Singhania RR. Cellulase adsorption on lignin: A roadblock for economic hydrolysis of biomass. Renew Energy. 2016;98:29–42.

39. Gomes D, Rodrigues AC, Domingues L, Gama M. Cellulase recycling in biorefineries—is it possible? Appl Microbiol Biotechnol. 2015;99:4131–4143.

40. Qing Q, Wyman CE. Hydrolysis of different chain length xylooliogmers by cellulase and hemicellulase. Bioresour Technol. 2011;102:1359–66.

41. Bai Flagfeldt D, Siewers V, Huang L, Nielsen J. Characterization of chromosomal integration sites for heterologous gene expression in *Saccharomyces cerevisiae*. Yeast. 2009;26:545–551.

42. Kaishima M, Ishii J, Matsuno T, Fukuda N, Kondo A. Expression of varied GFPs in Saccharomyces cerevisiae: codon optimization yields stronger than expected expression and fluorescence intensity. Sci Rep. 2016;6:35932–35932.

43. Inokuma K, Hasunuma T, Kondo A. Efficient yeast cell-surface display of exo- and endo-cellulase using the SED1 anchoring region and its original promoter. Biotechnol Biofuels. 2014;7:8.

44. Inokuma K, Kurono H, den Haan R, van Zyl WH, Hasunuma T, Kondo A. Novel strategy for anchorage position control of GPI-attached proteins in the yeast cell wall using different GPI-anchoring domains. Metab Eng. 2020;57:110–117.

45. Chen DC, Yang BC, Kuo TT. One-step transformation of yeast in stationary phase. Curr Genet. 1992;21:83–4.

46. Fang F, Salmon K, Shen MWY, Aeling KA, Ito E, Irwin B, et al. A vector set for systematic metabolic engineering in *Saccharomyces cerevisiae*. Yeast. 2011;28:123–136.

47. Guirimand G, Inokuma K, Bamba T, Matsuda M, Morita K, Sasaki K, et al. Cell-surface display technology and metabolic engineering of *Saccharomyces cerevisiae* for enhancing xylitol production from woody biomass. Green Chem. 2019;21:1795–1808.

48. Miller GL. Use of dinitrosalicylic acid reagent for determination of reducing sugar. Anal Chem. 1959;31:426–428.

49. Basso LC, de Amorim HV, de Oliveira AJ, Lopes ML. Yeast selection for fuel ethanol production in Brazil. FEMS Yeast Res. 2008;8:1155–63.

50. Schwan RF, Mendonca AT, da Silva JJ, Rodrigues V, Wheals AE. Microbiology and physiology of Cachaca (Aguardente) fermentations. Anton Leeuw Int J G. 2001;79:89–96.

