## Additional file 1 for "Consolidated bioprocessing of corn cob-derived hemicellulose: engineered industrial *Saccharomyces cerevisiae* as efficient whole cell biocatalysts"

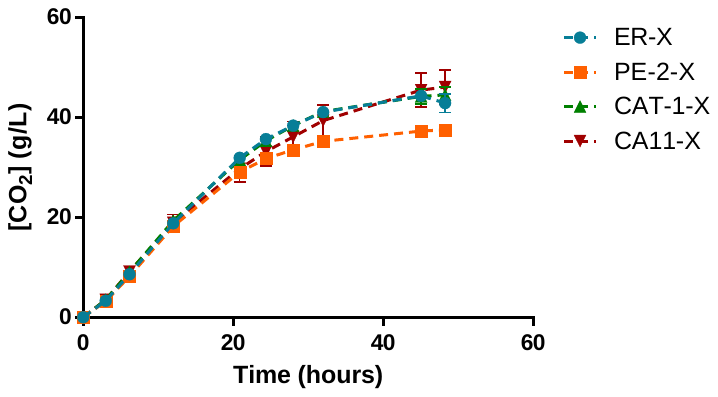

**Fig. S1**. Fermentation in corn cob liquor 29X_Pot_ (75 %) supplemented with 100 g/L glucose at 40 °C to evaluate the metabolic capacity of the different strains at higher temperature. Cells were inoculated at 10 g wet cells/L in flasks fitted with glycerol lock. The fermentation was performed at 40 °C and 150 rpm for 48 hours and was monitored by measuring the reduction of mass loss resulting from CO_2_ production. Data represents the average ± SD from two biological replicates.

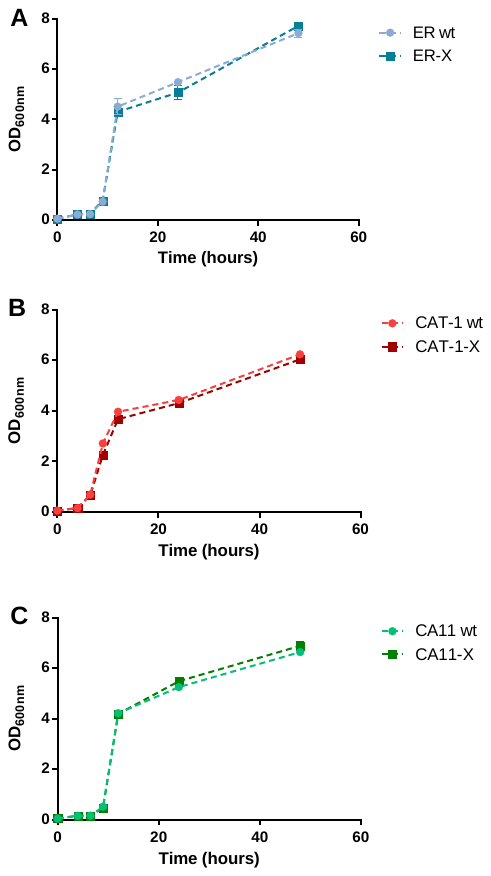

**Fig. S2**. Growth profiles of the wild type *S. cerevisiae* strains, ER, CAT-1 and CA11, and of the strains with cell surface display of hemicellulolytic enzymes, ER-X, CAT-1-X and CA11-X, in YPD. Cells were inoculated at an OD_600nm_ of 0.05 and grown for 48 hours at 30 °C and 200 rpm. Data represents the average ± SD from two biological replicates.

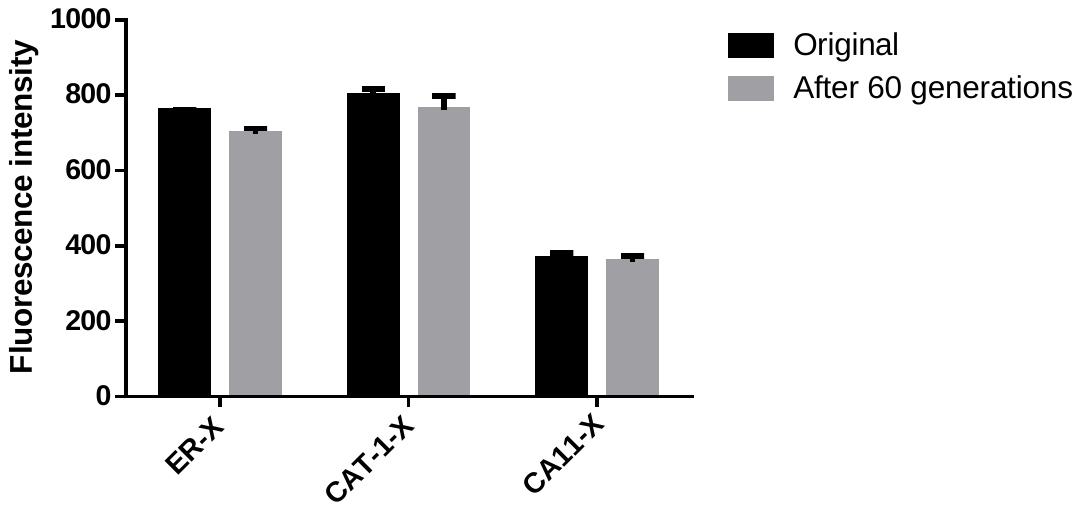

**Fig. S3**. Stability evaluation of the integration at the δ-sequences after growth for 60 generations. Cells were grown in YPD liquid medium for 60 generations. The fluorescence intensity of the cells before and after 60 generations (normalized to an OD_600nm_ of 0.2) was measured in a microplate fluorometer set at 480 nm (excitation) and 510 nm (emission). Data represents the average ± SD from two biological replicates.

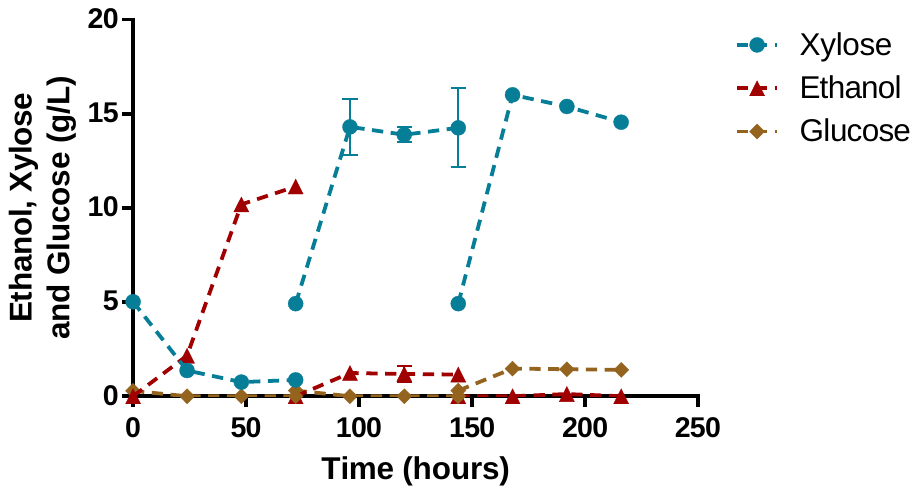

**Fig. S4**. Fermentation in corn cob liquor 32X_Pot_ with ER-X-2P strain and 2 cycles of cell recycling. Data represents the average ± SD from two biological replicates.

**Table S1**. β-glucosidase 1 (BGL1), β-xylosidase A (XYLA) and endoxylanase II (XYN) enzymatic activities.

| Strain | BGL1 (U/g DCW) | XYLA (U/g DCW) | XYN (U/g DCW) | |
| --- | --- | --- | --- | --- |
|  |  |  | 30 °C | 40 °C |
| ER | 39.1 | 87.8 | 139 ± 24 | 263 ± 2 |
| PE-2 | 71.1 | 155 | 93.1 ± 2.2 | 162 ± 1 |
| CAT-1 | 50.6 | 50.2 | 76.4 ± 3.7 | 140 ± 2 |
| CA11 | 71.3 | 47.8 | 41.0 ± 8.0 | 57.9 ± 1.1 |

**Table S2**. Plasmids and primers used for subcloning steps in this study. Lower case sequences indicate addition of homologous regions for plasmid assembling.

| **Primers** | **Sequence (5’->3’)** | **Aim** |
| --- | --- | --- |
| pAll_fw | CGCATCAGGAAATTGTAAACG | Amplification of pIBG-SSA for construction of pI23-BGL1-kanMX; Amplification of pIU5-TeCBH1c-SSS for construction of pI5-CBH1-NatMX |
| pBGL1_CBH2_rv | CGTCAGGTGGCACTTTTC | Amplification of pIBG-SSA for construction of pI23-BGL1-kanMX |
| I23-BGL1_fw | aagtgccacctgacgACAGAGAAGGACAAGGCTGAAG | Amplification of intergenic region 23 from ER for construction of pI23-BGL1-kanMX |
| I23-BGL1_rv | cgacctgcagcgtacGACACAGGTGACAATAAAGTTTCC |  |
| kanMX-BGL1_fw | GTACGCTGCAGGTCGACAAC | Amplification of *kanMX* resistance marker for construction of pI23-BGL1-kanMX |
| kanMX-BGL1_rv | caatttcctgatgcgATAGGCCACTAGTGGATCTGATATC |  |
| I23_FW | ATAATGAGTTCCGAGTCTGTTGGTG | Confirm integration in intergenic region 23 |
| I23_RV | CGAGATAAGGCATGGGGTTCTG |  |
| pI5-CBH1_rv | CAGGTTGTGCTCACTGTATATAGTC | Amplification of pIU5-TeCBH1c-SSS for construction of pI5-CBH1-NatMX |
| Nat-pI5-CBH1_fw | agtgagcacaacctgGACATGGAGGCCCAGAATAC | Amplification of nat*MX* resistance marker for construction of pI5-CBH1-NatMX |
| Nat-pI5-CBH1_rv | caatttcctgatgcgCAGTATAGCGACCAGCATTC |  |
| pI5-NatMX_fw | TGCTGGCGTTTTTCCATAG | Amplification of pI5-CBH1-NatMX for construction of pI5-XylA-NatMX |
| pI5-NatMX_rv | CAGCCTGAATGGCGAATG |  |
| XylA_fw | tcgccattcaggctgCTTCGCTATTACGCCAGATTG | Amplification of pIK-BX-SSS for construction of pI5-XylA-NatMX |
| XylA_rv | ggaaaaacgccagcaCGAATTGGGTACCTTTGATTATG |  |
| H-73 | TCTCTCTTGCACCAGCCATT | Confirm integration in intergenic region I5 |
| H-75 | CGGAATCGCATCAGGTCTT |  |
| pδW_fw | CTGAGAAATGGGTGAATGTTGAG | Amplification of pδW-EX-SSS for construction of pδW-XYN-kanMX |
| pδW_rv | CGGGGGATCCACTAGTTCTAG |  |
| kanMX-UkG1_fw | ctagtggatcccccgGACACACAAAATATCCCTTCCTACTC | Amplification of kanMX-UkG1 cassette for construction of pδW-XYN-kanMX |
| kanMX-UkG1_rv | tcacccatttctcagGGTGTCGACAACCCTTAATATAACTTC |  |
| pCRE_fw | CCATCTTGCACTTCAATAGCATATC | Amplification of pBF3060 for construction of pCRE-hyg6 |
| pCRE_rv | GTATGAGTATTCAACATTTCCGTGTC |  |
| hyg6_fw | gttgaatactcatacGACATGGAGGCCCAGAATAC | Amplification of *hyg6* resistance marker for construction of pCRE-hyg6 |
| hyg6_rv | tgaagtgcaagatggCAGTATAGCGACCAGCATTCAC |  |
| **Plasmids** | **Relevant features** | **Source** |
| pIBG-SSA | SED1p–SED1ss–*A. aculeatus* BGL1–SAG1a–SAG1t, *HIS3* | [1] |
| pIU5-TeCBH1c-SSS | SED1p–SED1ss–T. emersonii CBHI–SAG1a–SAG1t, *URA3* | [2] |
| pI5-CBH1-NatMX | SED1p–SED1ss–T. emersonii CBHI–SAG1a–SAG1t, *NatMX* | This work |
| pIK-BX-SSS | SED1p–SED1ss–*A. oryzae* XYLA–SED1a–SAG1t, *LYS2* | [3] |
| pδW-EX-SSS | SED1p–SED1ss–*T. reesei* XYNII–SED1a–SAG1t, *TRP1*, δ-integration | [3] |
| pBF3060 | CRE recombinase, *URA3* | [4] |

**References**

1.	Inokuma K, Hasunuma T, Kondo, A. Efficient yeast cell-surface display of exo- and endo-cellulase using the SED1 anchoring region and its original promoter*.* Biotechnol Biofuels. 2014;7:8.

2.	Liu Z, Inokuma K, Ho S-H, Haan Rd, Hasunuma T, van Zyl WH, et al. Combined cell-surface display- and secretion-based strategies for production of cellulosic ethanol with *Saccharomyces cerevisiae.* Biotechnol Biofuels. 2015;8:162.

3.	Guirimand G, Inokuma K, Bamba T, Matsuda M, Morita K, Sasaki K, et al. Cell-surface display technology and metabolic engineering of *Saccharomyces cerevisiae* for enhancing xylitol production from woody biomass*.* Green Chem. 2019;21:1795-808.

4.	Fang F, Salmon K, Shen MWY, Aeling KA, Ito E, Irwin B, et al. A vector set for systematic metabolic engineering in *Saccharomyces cerevisiae.* Yeast. 2011;28:123-36.
